# Bilateral retinofugal pathfinding impairments limit behavioral compensation in near-congenital one-eyed *Xenopus laevis*

**DOI:** 10.1101/2023.05.05.539408

**Authors:** Michael Forsthofer, Clayton Gordy, Meghna Kolluri, Hans Straka

## Abstract

To generate a coherent visual percept, information from both eyes must be appropriately transmitted into the brain, where binocular integration forms the substrate for visuomotor behaviors. To establish the anatomical substrate for binocular integration, the presence of bilateral eyes and interaction of both optic nerves during retinotectal development play a key role. However, the extent to which embryonic monocularly derived visual circuits can convey visuomotor behaviors is unknown. In this study, we assessed the retinotectal anatomy and visuomotor performance of embryonically generated one-eyed tadpoles. In one-eyed animals, the axons of retinal ganglion cells from the singular remaining eye exhibited striking irregularities in their central projections in the brain, generating a non-canonical ipsilateral retinotectal projection. This data is indicative of impaired pathfinding abilities. We further show that these novel projections are correlated with an impairment of behavioral compensation for the loss of one eye.

## Introduction

Integration of bilateral sensory information is crucial for detailed and environmentally relevant processing of the world. Such processing aids in the extraction of behaviorally important sensory features. In the visual system for example, vision with two eyes, or stereopsis, contributes to distance estimation^1^ and motion detection^2^. In order to accomplish this, anatomical pathways conveying separate information from the left and right eye must be integrated in the central nervous system to form a coherent visual percept^3^.

In some animals, such as mice, this combination of bilateral visual information happens at the level of retinal ganglion cell (RGC) projections: axons project into the brain either ipsi-or contra-laterally depending on their topographical origin in the retina. In other animals, such as zebrafish or larval *Xenopus* tadpoles, all axons cross to the contralateral side (for review, see ^4^). In this latter case, binocular information must still converge, which occurs anatomically via interhemispheric commissures, to form a binocular visual percept and facilitate binocular-dependent behaviors^5-8^.

Following loss or obstruction of one eye in mature organisms, these established circuits display an impressive capacity to compensate for loss in input, on both a functional and behavioral level. This even includes recovery of visual associated behaviors to certain extents^9-12^. In frogs, for example, acute monocular deprivation due to eyelid suture normally leads to direction-selective impairment of the optokinetic reflex (OKR). However, within 8 days this impairment is remedied, and the OKR responds equally to all directions even under monocular stimulation conditions^13^. To enable this plasticity in mature stages, circuits must be correctly assembled during embryonic development^14^. Such canonical assembly occurs with contribution from both eyes, with genetic and molecular factors playing a key role for pathfinding of bilateral RGCs, and neural activity serving to fine tune morphological arborizations and other aspects of neuronal connectivity^15-19^. Genetically induced deviations in chiasmatic crossing have a direct impact on visual behaviors, such as the complete ipsilateral RGC projections in the zebrafish *belladonna* mutant that cause a left-right inversion of eye-following motion in the optokinetic reflex^14,20^.

A large body of research was conducted on the genetic basis of the establishment of retinotectal connections and their specific functions. Less is known about the role of molecular or functional interactions of bilateral visual pathways during embryogenesis, and how these contribute to establishment of visual circuits and their functional output. In animals where each eye possesses native RGC projections to both the ipsi- and contralateral brain hemispheres, such as ferrets and mice, early embryonic ablation of an optic vesicle and subsequent one-eyed embryonic development results in reduced ipsilateral projections from the remaining eye^21,22^. This suggests that binocular RGC axons are involved in the chiasmatic-crossing decision, specifically in contributing to the ipsilateral projection of the contralateral eye.

In animals with exclusively contralateral RGC projections, similar experiments yielded less unanimous results. In chick embryos, removal of one optic vesicle induces an ipsilateral RGC projection from the remaining eye. However, these are lost within few days of development before any visual behavior in this species, and do not allow conclusions about anatomical or functional impact on post-embryonic behavior^23^. Unilateral embryonic eye removals in *Xenopus* on the other hand show no sign of an aberrant ipsilateral projection following metamorphosis^24^. However, these optic vesicle removals were performed after onset of RGC differentiation, and mammalian studies showed a large variability in phenotypes depending on timing of optic vesicle removal^25^. Additionally, neither study assessed function of circuitry formed under embryonic monocular conditions, leaving a potential beneficial or detrimental role of embryonic plasticity open. While a detrimental effect poses the more likely outcome based on previous genetic studies on pathfinding errors^14^, the complete absence of visual input to half of the circuit, rather than miswiring to both, may allow for stronger functional connections of the remaining eye, and to functional plasticity^26^. Indeed, *Xenopus* embryos show a large degree of plasticity during embryonic development, allowing for anatomical integration of e.g. a third eye into midbrain circuits^27^. More recent studies reveal that supernumerary or displaced eyes and ears can be anatomically and functionally connected to their central targets and facilitate behavior to certain extents and under certain conditions^28-31^.

In this study we revisit embryonic visual circuit formation in one-eyed *Xenopus laevis*, to answer the question whether visual circuits can develop appropriately in absence of one eye during embryogenesis, and whether this developmental plasticity conveys a compensatory or detrimental effect on visuomotor behavior. In contrast to previous studies in this species, we performed embryonic enucleations before differentiation of RGCs^16^. Following development into late larval stages, we expect exclusively contralateral RGC projections in two-eyed animals. In monocular animals, we unexpectedly found an ipsilateral RGC projection. We then quantified eye motion elicited by the optokinetic reflex, to assess the extent of functional plasticity of this monocular visual circuit. The results reveal that embryonically generated one-eyed *Xenopus* tadpoles with aberrant RGC projections exhibit more impaired visuomotor behaviors, highlighting a critical limitation of visuomotor plasticity in retinotectal development.

## Results

### Generation of one-eyed tadpoles

One-eyed tadpoles were generated by removal of the left optic vesicle in early embryonic stages, similar to embryonic removal of the otic placode in previous studies (Fig. 1a, Stages 26-28)^32,33^. Removals were validated one week post-surgery at stage 46 by visual inspection (Fig. 1b, top, Suppl. fig. 1a, b, 26.5 % success rate) in comparison with unmanipulated controls (Fig. 1b, bottom, Suppl. fig. 1c). To further visualize the observed absence of gross anatomical optic structures, immunohistochemical labelling of cranial nerves (acetylated tubulin, green) and extraocular musculature (myosin VI, red) was performed. On the unmanipulated side in these animals (Fig. 1a, left inset), and in unmanipulated controls (Fig. 1a, right inset), the extraocular muscles insert themselves in a stereotyped fashion around the eye and are innervated by their respective extraocular motoneurons (Fig. 1a, right inset), consistent with features for *Xenopus laevis*^34^. In embryonic one-eyed animals however, such an organizational scheme was not observed (Fig. 1a, left inset), given the absence of an eye in this region. Lacking individual eye muscles to innervate, the suspected extraocular motor neurons in the optic-area of these animals, identified based on their exit sites from the skull, failed to innervate in canonical fashion (Fig. 1a, insets, arrowheads). This immunohistochemical data validated the absence of eye associated neural and muscular structures in embryonically manipulated animals^35^. Monocular animals were then further reared until stages 52-54^32^, where visuomotor behaviors and central retinotectal circuits are readily profiled^36^, in order to anatomically and behaviorally asses the effects of embryonic removal of the optic vesicle. *In-vitro* preparations (Fig. 1a, right, Suppl. fig. 1 d, e) of control and one-eyed animals were generated and subsequently subjected to functional assessment paradigms which have been previously shown to induce quantifiable optokinetic eye movements (Fig. 1c; ^37^).

**Figure 1:**
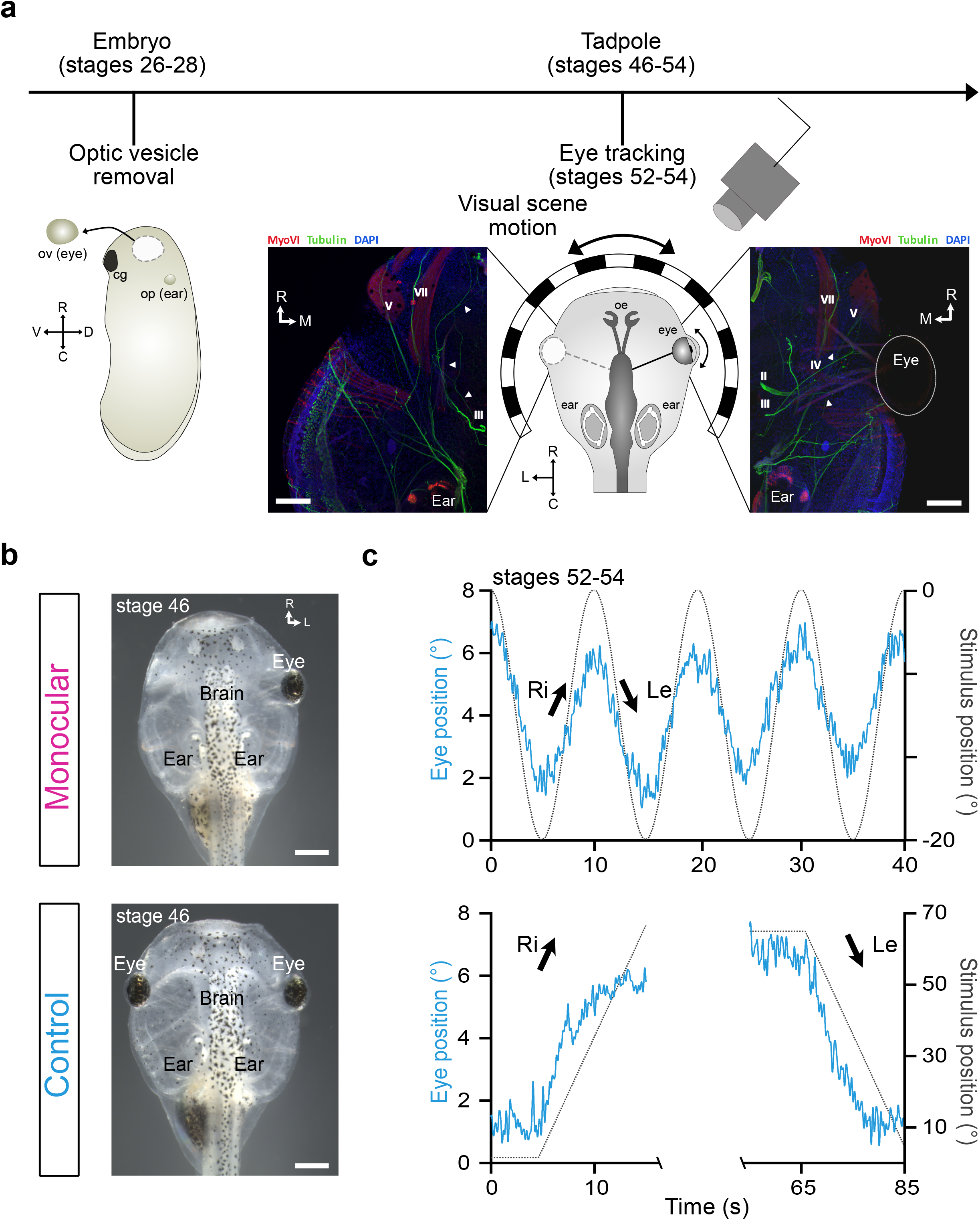
Methodology of eye removals and behavioral tracking. a) Schematic of the experimental timeline, consisting of embryonic removal of the left optic vesicle at stages 26-28 (left, total of 83 animals, 22 successful) and behavioral tracking in a visual stimulation paradigm following maturation at stages 52-54 (right). Insets depict representative immunohistochemical staining of the eye region in a one-eyed (left) and a control (right) tadpole at stage 46, showing nerves (Tubulin, green), muscles (MyoVI, red) and a nuclear counter-stain (DAPI, blue). Arrows indicate the trajectory of the oculomotor nerve. b) Representative images of one-week old one-eyed (top) tadpoles in comparison to controls (bottom). c) Representative traces of right eye motion in response to sinusoidal motion stimulation (top, 6,28 °/s peak velocity at 0.1 Hz) or constant velocity unidirectional motion (bottom, 6.28 °/s) in a single control animal. Eye motion depicted in blue, stimulus motion in black. Scalebars: a) 200 μm, b) 500 μm

### OKR performance in bidirectional stimulation

In control animals with two bilateral eyes (hereafter, “binocular”), visuomotor behaviors were profiled following visual stimulation. For this, we presented a sinusoidal horizontally moving black-white striped pattern, which readily induced reflexive eye movements that followed the motion cyclically (Fig. 1a, c, Fig. 2a). In binocularly stimulated controls, eye motions were conjugated between both eyes (Suppl. fig. 2a, b, slope = 0.92 ± 0.01, mean ± SD) and showed a gain of 0.28 ± 0.11 (Fig. 2a, b, blue, eye motion amplitude/stimulus motion amplitude) in line with previous studies in this species^33^. To better understand the individual contributions of each eye to these bilateral control visuomotor behaviors, we next investigated OKR performance following acute loss of visual input on one side. This was induced by selective lesioning of the left optic nerve (hereafter, “Lesioned”) close to the brain as previously described for the statoacoustic nerve in such preparations^37^, which induced an acute monocular condition. Such a condition allowed us to assess the extent to which mature and entrained visuomotor circuits are affected by sudden loss of input from one side. In lesioned animals, the OKR gain decreased to 0.18 ± 0.05 (Fig. 2a, b, green, p=0.041, Kruskal-Wallis test with Dunn’s multiple comparison), indicating an impairment after acute unilateral eye loss. Coverage of the left visual field instead of surgical intervention yielded the same effect of a decrease of the OKR gain from 0.34 ± 0.06 to 0.21 ± 0.04 (Data not visualized, p=0.015, Wilcoxon signed rank test). As impairments from surgical lesions and restrictions of the visual field yielded similar results, the OKR impairment likely originated from visual signal loss rather than detrimental effects of the surgical procedure itself.

**Figure 2:**
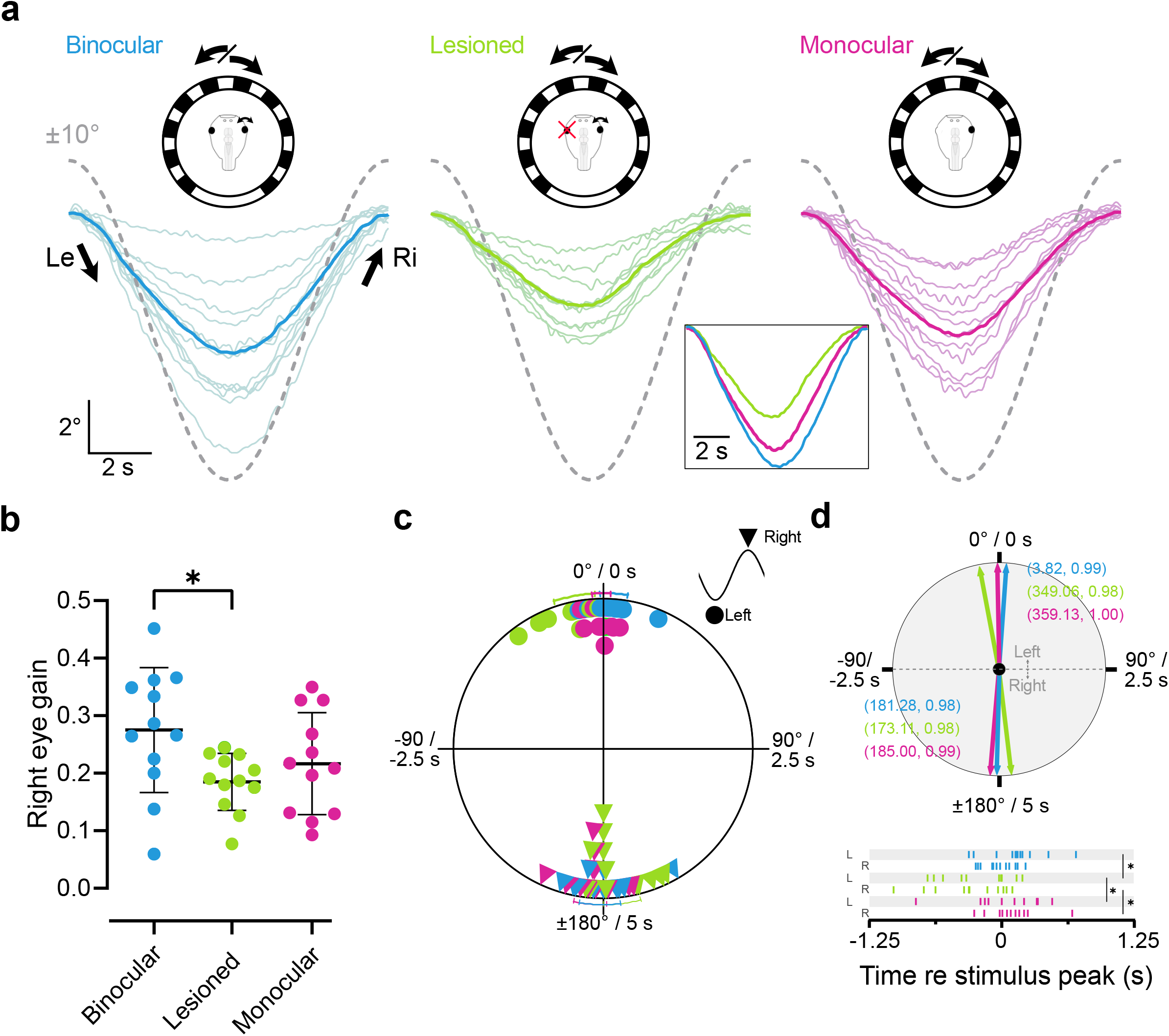
Responses of the right eye to sinusoidal motion stimulation. a) Eye motion traces (colored lines) of the right eye of binocular (n = 12), left eye lesioned (n = 12) and monocular tadpoles (n = 12) in response to sinusoidal stimulus motion (dotted lines). Thin lines represent average responses per animal (8-30 cycles per animal), thick lines in plots and in the inset represent the average response across animals. Schematics depict stimulation paradigm. b) Gain values (eye motion amplitude/stimulus amplitude) for the binocular, lesioned and monocular paradigms. c) Average phase values of the eye following motion respective stimulus motion in degrees and seconds. Data points represent animals, with standard deviation bars outside the circle. 0° represents the peak at the leftward to rightward direction change, 180° the reverse. d) Top: Mean vectors (length, vector strength) of the data plotted in c) bottom: timing values are given in relation to the stimulus motion peak corresponding to the leftward or rightward eye motion peak. Data re-used from panel c. L: left to right peak, R: right to left peak.

With reference values for OKR performance in binocular and left eye lesioned tadpoles, we next examined whether embryonically one-eyed tadpoles (Fig. 1a, b, Suppl. fig. 1b, d; hereafter, ‘monocular’) were able to compensate for impairment of the OKR in monocular stimulation conditions. These tadpoles displayed eye movements in response to sinusoidal stimuli at a gain of 0.22 ± 0.09, which placed them in between binocular and lesioned tadpole performances, with no significant difference to either group (Fig. 2a, b, magenta, p = 0.36 and p>0.99, respectively, Kruskal-Wallis test with Dunn’s multiple comparison). Accordingly, the OKR in these animals was not strongly impaired during exclusive right eye stimulation and suggests a degree of compensation for the lack of one eye.

In acute optic nerve lesions, a change in the timing of the OKR response peak from a mostly in-phase response for both leftward and rightward motion (Fig. 2c, d, blue, 3.8 ± 8.0°, 1.3° ± 11.7°, respectively) to a phase lead in the leftward (Fig 2c, d, upper half, green, -11.0 ± 11.8°, p=0.002, Watson-Williams F-test) but not the rightward direction (Fig 2c, d, lower half, -7.9±10.8°, p = 0.1, Watson-Williams F-test, Fig. 2c, bottom, d), and indicate that not just response strength, but also timing of OKR is impaired upon acute loss of one eye. Response latencies of embryonic one-eyed animals were clearly improved from lesioned controls in both leftward and rightward direction (Fig. 2d, magenta, -0.97 ± 5.3°, p = 0.017, 5 ± 9.5°, p = 0.012, Watson-Williams F-test) and were similar to binocular animals (p = 0.117, p = 0.103, Watson-Williams F-test).

Overall, sinusoidal visuomotor stimulation reveals that tadpole OKR requires binocular visual input to maintain strength and timing of the response, and loss of one eye impairs both factors. Tadpoles which only developed with one eye, however, were able to partially compensate for these impairments.

### Direction-specific impairment of OKR

As our manipulation induced an asymmetry on bilateral peripheral sensory detection, we next investigated if the impairment also affected the OKR in a direction specific, and therefore asymmetric manner. In many lateral eyed animals, each eye in isolation preferentially responds to motion in the nasal (N) direction^38^. We therefore tested whether the amplitude impairment observed in sinusoidal stimulation was caused by a reduction of the temporal (T) motion component, in other words, a rightward movement in case of the singular right eye. To assess this, we measured eye velocity in the first two seconds during unidirectional motion stimulation at a constant velocity. In binocular control tadpoles, during binocular stimulation, the right eye responded more strongly to temporal motion at a gain of 0.39 ± 0.12 (eye velocity/stimulus velocity) versus 0.24 ± 0.07 for nasal motion (Fig. 3a, b, blue, p<0.001, two-way ANOVA, Bonferroni’s multiple comparison). Therefore, under binocular viewing conditions, the right eye responds well in both motion directions with a preference for temporal motion. This temporal preference was present in both eyes, as confirmed by comparative conjugation assessment (Suppl. Fig. 2a), and only in unidirectional but not sinusoidal stimulation (Suppl. Fig. 2b). This indicates a control condition whereby the OKR exhibits an asymmetry which favors temporal motion detection.

**Figure 3:**
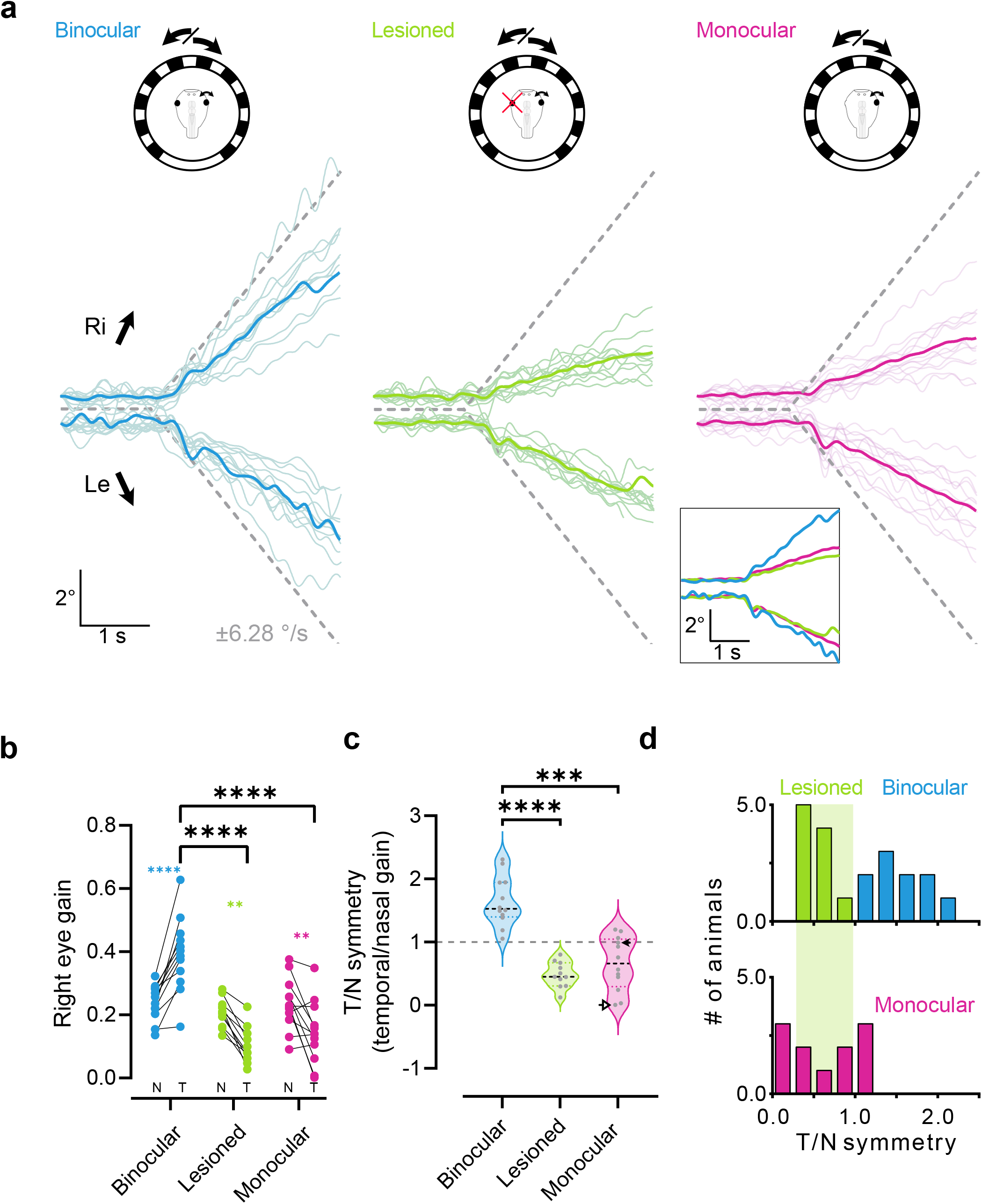
Responses of the right eye to unidirectional motion stimulation. a) Motion traces (colored lines) of the right eye of binocular (n = 12), left eye lesioned (n = 11) and monocular tadpoles (n = 12) in response to unidirectional stimulus motion (dotted lines). Thin lines represent average responses per animal (1-4 stimulus bouts per direction), thick lines in plots and in the inset represent the average response across animals. b) Gain values (eye velocity/stimulus velocity) within the first two seconds of motion onset for the three paradigms. Connected dots represent nasal (N) and temporal (T) motion of the same eye. c) Symmetry indices (temporal gain/nasal gain) for the three paradigms. Values >1 indicate stronger motion towards the right, <1 motion towards the left. 1 (grey dotted line) indicates equal motion to the left and the right and a symmetric response. Arrowheads indicate low and high representative datapoints corresponding to anatomical values in Fig. 4e) and comparisons of anatomy and function in Fig. 4f) d) Histograms of the distribution of symmetries in binocular and lesioned (top) versus monocular (bottom) animals. Data taken from panel c). Bin width on the x-axis is 0.25. Green shading highlights the distribution of lesioned animals for easier comparison with monocular animals.

Acute optic nerve transection of one eye selectively impaired OKR of the remaining eye in the temporal direction. Under such monocular viewing conditions, gain of the right eye in the temporal direction was reduced to 0.10 ± 0.05 and therefore lower than in binocular tadpoles (Fig. 3b, green, p<0.001, two-way ANOVA with Bonferroni’s multiple comparison), while nasal motion was unaffected at a gain of 0.19 ± 0.05. Therefore, under monocular conditions, the remaining eye shows a direction preference for nasal stimulus motion (Fig. 3b, green, p<0.001, two-way ANOVA, Bonferroni’s multiple comparison, Fig. 3c, 0.48 ± 0.2°, p<0.001, Kruskal-Wallis test with Dunn’s multiple comparison), in line with other lateral-eyed animals^38^. We hypothesize that this is due to a cooperative processing of temporal motion of one eye with nasal motion of the other one, as an addition of nasal and temporal gain yields similar gain values as temporal motion in binocular conditions. Overall, these lesions confirm that acutely induced asymmetry in the visual sensory periphery imposes an asymmetry on visuo-motor transformation.

Having established a direction-specific motor impairment of unilateral loss of visual input in post-embryonic tadpoles, we next tested whether tadpoles that were reared from early embryo stages with only one eye compensated for such a sensory asymmetry during development. In contrast with the compensation shown by monocular tadpoles during sinusoidal stimulation, temporal eye motion during unidirectional constant velocity stimulation exhibited a gain of 0.14 ± 0.1 which was still impaired compared to binocular animals (p < 0.001, two-way ANOVA, Bonferroni’s multiple comparison). Rather, this gain level was comparable to animals with an acute loss of one optic nerve (Fig. 3a, b, magenta). Moreover, motion in the nasal direction was not different than controls at 0.24 ± 0.08 (p = 0.58, two-way ANOVA, Bonferroni’s multiple comparison), and as a result, the OKR was asymmetric towards the nasal direction (Fig. 3c, magenta, 0.65 ± 0.42, p=0.0007, Kruskal-Wallis test, Dunn’s multiple comparison). Therefore, on a population level embryonic one-eyed animals appear to not exhibit behavioural plasticity sufficient to overcome an impairment in OKR directional perception. However, within this cohort of embryonic monocular tadpoles, left-right asymmetries were more heterogeneous in comparison to lesioned tadpoles. Indeed, the symmetry index (temporal gain / nasal gain, Fig. 3c) of monocular tadpoles did not distribute in a Gaussian fashion (Fig. 3c, d, magenta, Kolmogorov-Smirnow test, p=0.0038. Data taken from c.), which is in contrast to both binocular and lesioned tadpoles (Fig. 3c, d, blue and green). This was likely caused by individual monocular tadpoles with an equal leftward and rightward, and therefore symmetric, OKR response (Fig. 3c, grey line, Fig. 3d, bottom). Such behavioral variability following embryonic unilateral sensory deprivation has been previously reported^33^, and we next investigated whether this functional variability had anatomical correlates following monocular embryonic development.

### Anatomical correlates - RGC projections

As the OKR in *Xenopus* is mediated by the pretectum^39,40^, a direct target of retinofugal fibers, we directly investigated the connectivity of RGC central projections to this area in binocular and monocular (embryonically manipulated) tadpoles. Application of dextran-conjugated dyes into the eyes of *in-vitro* preparations (Fig. 4a, b, insets) revealed differences in RGC axon projections between one-eyed and control tadpoles. Control tadpoles showed exclusively contralateral RGC projections, reaching targets in the pretectum optic tectum (Fig. 4a, c) and thalamus (Suppl. fig 2f, left)^41,42^. In monocular tadpoles, a subset of RGC axons innervated ipsilateral thalamic, pretectal, and tectal targets (Fig. 4b, d, Suppl. fig. 2f, right) instead of, or in addition to, contralateral ones. We quantified the number of labelled processes in the anterior midbrain across the medio-lateral aspect of the brain (Fig. 4c, d) for multiple animals (controls: n=5, monocular: n=9), to approximate the amount of erroneously non-crossing RGCs (Fig. 4e). While in both the control and experimental conditions, most projections terminated in the contralateral anterior tectum (Binocular: 98.74 ± 0.4%, Mean ± SD, Monocular: 83 ± 21.6%, Mean ± SD), only monocular animals showed a fraction of RGC projections to the ipsilateral anterior tectum (Fig. 4e, f, 16.82 ± 21.62%, mean ± SD, p = 0.03, two-tailed Mann-Whitney test, Data in 4f taken from 4e). The number of ipsilateral projections was variable and heterogenous within the monocular group, particularly compared to binocular controls (Fig. 4e, f). Overall, we found that RGC pathfinding in one-eyed tadpoles is mainly impaired at the optic chiasm, with fibers projecting into the ipsilateral or contralateral pretectum and tectum. Further, the extent of chiasmatic pathfinding errors is highly variable between individuals, and reminiscent of the behavioral variability found within monocular tadpoles.

**Figure 4:**
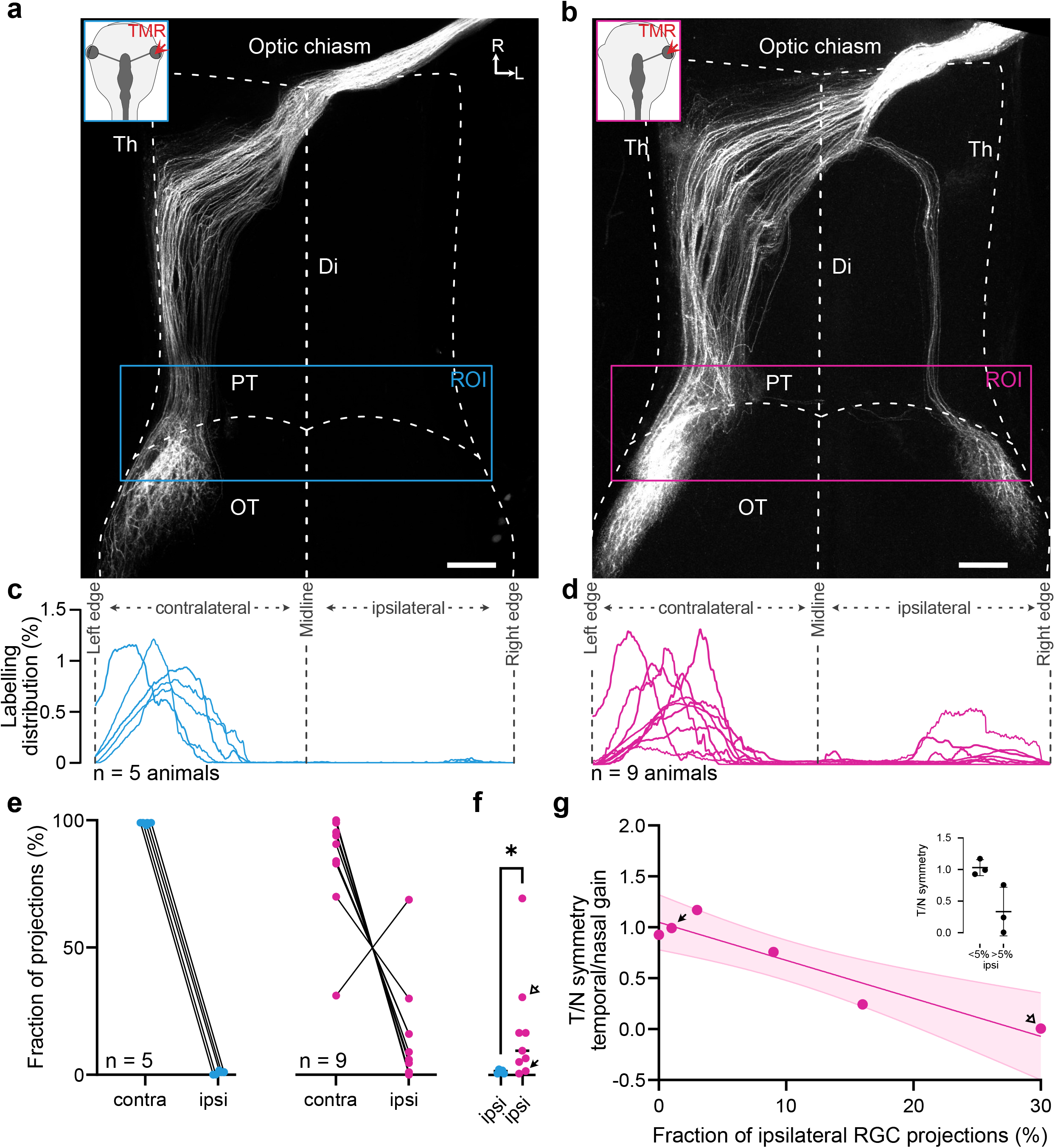
RGC projections in control and one-eyed tadpoles at stages 52-54. Representative confocal z-stacks of cleared whole brains of one control (a) and monocular (b) tadpole with RGC axons labelled. White background insets indicate dye application site (colored arrowheads), colored boxes indicate the region of interest used for quantification of projections. c, d) Histograms of the amount of above threshold labelling within ROIs in a), b) across the medio-lateral axis of the brain at the anterior tectum for all control (n=5) and monocular (n=9) preparations. e) Cumulative sum per animal of the sections of the histograms corresponding to the contralateral or ipsilateral part of the ROI in control (blue) and monocular (magenta) animals. f) Statistical comparison of the amount of ipsilateral signal in control and monocular animals. Arrows indicate datapoints corresponding to two representative datapoints for low and high ipsilateral projection values as found in Panel g) and Fig. 3c). g) Correlation of the fraction of ipsilateral RGC projections and symmetry of OKR in animals with anatomical and behavioral data (n=6). Negative slope indicates a negative correlation between the amount of ipsilateral RGC projections and OKR symmetry. R^2^ = 0.884, fit: y = -3.734y + 1.049. Inset shows comparison of symmetry values between two groups of animals, grouped based on an anatomical threshold (>5% ipsilateral projections). Th Thalamus, PT Pretectum, OT Optic tectum, Di Diencephalon, OT Optic Tectum, R rostral, L lateral, ROI Region Of Interest. Scalebars = 200 μm

We thus asked if this variability correlated to the previously described variability in OKR behavior. We compared anatomical and behavioural data in animals in which anatomical dye tracings were successfully performed following behavioural assessment (n=6). Plotting of the fraction of ipsilateral projections *versus* the symmetry index of the OKR response was used to approximate such a correlation. Linear fits through this relation revealed a significant negative correlation (Fig. 4g, red, r^2^ = 0.88, p = 0.005 compared to a slope of 0). Representative high (filled arrow) and low (empty arrow) OKR symmetry index animals (marked through Figs. 3c, 4f) suggest that an increased amount of erroneous ipsilateral RGC projections had a detrimental effect on functional compensation of the OKR. As nasal motion of these animals is unimpaired (Fig. 3b) and does not correlate with ipsilateral fibers (Suppl. fig. 2e, light blue) whereas the temporal direction does (Suppl. fig. 2e, red), it is likely that miswiring-caused impairments are either selective for temporal motion or can be compensated for in the nasal direction. Accordingly, animals with mostly or exclusively contralateral RGC projections partially compensated for the temporal OKR impairment following loss of one eye. While this is not a full restoration to the temporally asymmetric responses of binocular control tadpoles, it provides evidence for compensation of the temporal impairment observed in acute left-eye lesioned tadpoles as observed in adult frogs^43^.

Overall, our data show that embryonic development of the visual system under monocular conditions causes aberrations in retinotectal pathfinding of the remaining eye. These anatomical aberrations directly correlate with functional impairment of OKR symmetry and an impairment of compensatory plasticity to a symmetric OKR, as shown by monocular tadpoles with little RGC pathfinding errors.

## Discussion

The results of this paper demonstrate the requirement of bilateral eyes for appropriate establishment of contralaterally projecting RGCs in *Xenopus* tadpoles. They further show the detrimental effect of monocularity-derived retino-pretectal pathfinding errors on compensatory plasticity of behavior.

### OKR circuit development

Our results reported on the influence of binocularity on complete RGC axon decussation at the optic chiasm during embryonic development of *Xenopus*. While induction of ipsilateral retinotectal fibers has been demonstrated in this species previously, such results were observed in adult animals and resulted from regenerative, rather than embryonic, axonal growth^44,45^. Due to different molecular environments in embryonic *versus* adult tissue^46^, this regenerative effect does not necessarily translate to embryonic axonal pathfinding^45^. In fact, studies on embryonic axonal pathfinding in *Xenopus* were thus far unable to show a role of binocular interaction to establish the exclusive contralateral retinofugal projection during embryo development^24^. The discrepancy of our data with this study can be explained by two differences in study design: First, extirpations in our study are performed earlier in embryonic development at stages 26-28, as opposed to stage 30 in previous studies. Since RGCs already start differentiating at stage 28 with their axons being detectable in the optic stalk^47^, our removals at earlier stages of optic development may be more effective at preventing induction of an environment at the optic chiasm or in the optic stalk that facilitates contralateral, or prevents ipsilateral guidance. Second, as anatomical data in the earlier study was gathered in adult frogs, it is possible that the ipsilateral fibers we observe in larval stages disappear during metamorphosis. Indeed, a transient, non-canonical ipsilateral RGC innervation in the absence of a contralateral eye has been shown previously in chick embryos^23^; however, aberrant projections are lost again during embryonic development in this species, before potential manifestation of behavioral relevance of this pathway. The role of the contralateral eye in RGC pathfinding is further corroborated by studies in mice and ferrets. These studies, however, show an opposite effect to ours, where rather than inducing a non-canonical ipsilateral projection, a canonical ipsilateral projection is reduced^21,22^. Overall, while the importance of bilateral eyes for facilitation of appropriate decussation at the optic chiasm has been known, our study is the first to our knowledge to show this in *Xenopus*, while also highlighting an interspecies difference in the role of the contralateral eye as an inhibitor, rather than a promoter of ipsilateral RGC guidance.

Our study suggests the existence of mechanisms in chiasmatic pathfinding that remain thus far unidentified. Molecularly, Ephrin B is implicated in the formation of ipsilaterally crossing fibers in this species^48^. In fact, adult *Xenopus* display ipsilaterally projecting RGC axons, which are established during metamorphosis and can be induced pre-metamorphically in the presence of Ephrin B^48,49^. However, these axons target the thalamus rather than the tectum, and our results suggest that other mechanisms are relevant for axonal crossing in this species, either of a chemical or mechanical nature^50,51^. Even though we are unable to verify whether individual fibers target the ipsilateral equivalent of their canonical contralateral target area, overall pathfinding to gross pretectal and tectal areas appears intact^52^.

### Directionality in Xenopus OKR

We showed that anatomical deviations from canonical, all-contralateral RGC fibers are negatively correlated with functional compensation for monocularity in the OKR. In our tadpoles, compensation manifests as an improvement of the temporal motion direction in some embryonically monocular tadpoles. Curiously, we also observe an asymmetry under control binocular conditions in the temporal direction. In zebrafish, such asymmetries are associated with higher spatial frequency of stimulus patterns than the ones employed in our study, suggesting a different spatial frequency preference in *Xenopus*^53^. The stronger temporal component could originate from a summation of a small, temporal contribution from the respective eye with a stronger, nasal contribution of the contralateral eye during initial motion detection. This speculation is based on the symmetry and gain values we obtained by summing the temporal and nasal gain under monocular stimulation (Suppl. fig. 2c, d, grey dots). This may be mediated via intratectal and pretectal commissural connections, as in zebrafish^54^.

Preference for nasal motion in monocular stimulation is expected and fits data from other species^38,54,55^. In our experiments, acute loss of one eye leads to an impaired OKR in the remaining eye as expected. When such acute condition responses were compared to embryonically generated one-eyed animals, we found that behavioral compensation was possible, but correlated with retinotectal connectivity phenotypes. Behavioral compensation for loss of one eye was observed predominantly in animals with near-canonical, contralateral retinofugal pathways, and less in animals with ipsilateral retino-pretectal fibers. Compensation, however, does not restore OKR back to binocular levels with a temporal motion preference, but rather increases temporal motion to be equivalent with the nasal component, to facilitate symmetric OKR responses. Functionally, this could have two possible reasons: 1) Reaching maximum temporal motion requires bilateral eyes, and plasticity mechanisms are insufficient to compensate for the loss of the entire visual input from one. Or, 2) while each eye shows asymmetric OKR for unidirectional motion in binocular animals, the left and right eye cooperate to ensure an equal leftward and rightward response across the entire visual system, comprised of bilateral, complementary eyes. In monocular animals with control-like projections of the remaining eye, the single eye represents the entire visual system, and the only way to facilitate a symmetric OKR response is a symmetric response of this singular eye. Therefore, symmetry of the overall system, rather than a maximized motor response, may be desirable. Multiple studies back up the second hypothesis. First, embryonically generated one-eared animals compensate in a similar fashion in vestibular processing: At the cost of the normally preferred ipsiversive direction, the normally weak contraversive motion detection is increased to facilitate bidirectional motion sensing^33^. Second, multiple studies suggest that homeostasis is a common occurrence in tadpole plasticity, rather than response maximisation. In both VOR and OKR, training stimuli induce motor output homeostasis rather than stronger compensation^56,57^. Theoretical studies corroborate this finding, and suggest that consistency may be preferred, and full compensation may not be necessary to minimize retinal image slip under naturalistic conditions^58^.

### Role of ipsilateral RGC fibers

As only animals with no major ipsilateral projection show OKR compensation, we speculate that while these connections are likely functional on a physiological level, they impair appropriate sensorimotor function rather than contributing to compensation. They thus provide a possible alternative explanation for the impairment for the normally preferred motion direction in the vestibular system of embryonically one-eared tadpoles^33^. The detrimental effect of ipsilateral retinotectal connections may stem from functional miswiring, as RGCs selective for one motion direction wire to brain areas geared to process motion in the opposite direction. In *Xenopus* experiments placing a supernumerary eye or ear onto the head or even the tail, sensory afferents are either capable of functionally integrating or wire directly to their sensory nuclei, in large part probably due to conserved molecular guidance cues^27,28,30^. Aside from erroneous crossing, the ipsi-or contralateral canonical pretectal and tectal targets of RGCs here are correctly innervated, and our study aggrees with the notion that retinotectal and pretectal wiring is based majorly on genetic and molecular factors^59-62^. Within target nuclei, in addition to such factors, activity plays a major role to establish map formation and appropriate connectivity to postsynaptic neurons^63^. In experiments in zebrafish and *Xenopus*, ipsilateral connections from the eye to the tectum were induced through ablation of one tectal hemisphere. In both cases, these rerouted neurons targeted their appropriate topographical targets within the tectum, and in case of zebrafish, even wired to the same directionally selective neurons as their ipsilateral counterparts along the rostro-caudal axis^42,64^. In absence of an appropriately wired eye like in the present study, such direction appropriate wiring is likely to be detrimental: temporally selective RGCs of the right eye respond to rightward stimulus motion. Appropriately wired fibers to contralateral temporally selective pretectal neurons in the left pretectum will then initiate a rightward OKR^39,65^. However, if contacted, ipsilateral pretectal cells in the right pretectum initiate a leftward OKR in response to temporal stimulation, which would normally come from a leftward motion detected by the left eye. These competing signals may impair restoration of symmetric OKR. Multiple studies show similar behavioral effects of erroneous RGC projections in zebrafish mutants, where completely inversed retinotectal projections lead to an inversion of OKR^14,20^. Likewise, inversely connecting the left and the right eye with their ipsilateral tectum in adult frogs through surgery achieves inversion of the OKR^66^ in adult frogs. It is hypothesized that temporal motion is conveyed by pretectal commissures^54,67^ while nasal motion stems from direct motor neuron activation^40^. Behavioral impairments following retino-pretectal miswiring in our study therefore may go hand in hand with errors in commissural signaling, or wrongly crossing fibers originating predominantly from temporal direction selective RGCs.

As changes in ipsi-or contralateral connectivity are associated with weaker compensation, adult plasticity mechanisms likely play a role in the observed compensation. In adult mice, functional plasticity in response to impairments in direction perception is following monocular deprivation^9,68^, as it is in frog^43^. In the latter case, evidence suggests that the pretectum itself plays a role in maintaining directional symmetry^43^. Another likely candidate in *Xenopus* is the cerebellum, which is in general associated with OKR plasticity^69^. Specifically the gain up-regulation required for the observed compensation in the temporal direction is linked to the cerebellum in *Xenopus*, which is close to functionally mature at our employed developmental stages^56^. Coupling of embryonic eye removals with developmentally early inactivation of the cerebellum could help identify contributions of the cerebellum either as a mediator or initiation site of such plasticity. Other brain areas are equally probable, particularly the hindbrain vestibular nuclei, which receive visual motion information and aid in consolidation of cerebellar-aided gaze stabilizing plasticity Such contributions could be elucidated by means of visuo-vestibular mismatch studies. Finally, the pretectum itself has been shown to be involved in facilitation or impairment OKR symmetry following lesion-based impairments and warrants further functional investigations^70-72^.

Overall, this study highlights two main aspects relating to plastic compensation of the OKR: In *Xenopus*, bilateral eyes are essential to reliably develop contralateral retinofugal fibers as previously shown in other species. Moreover, aberrant ipsilateral retino-pretectal connections do not appropriately mediate visuomotor behaviors in absence of another eye. Rather, they are correlated with impairments of specifically temporal eye motion, suggesting that embryonic pathfinding changes put a limit on optokinetic plasticity.

## Acknowledgements

The authors would like to thank Prof. Dr. Ruben Portugues for support in finalizing and submitting the manuscript after the unexpected passing of Prof. Dr. Hans Straka in December 2023. The authors would like to thank Dr. Francois Lambert, Prof. Boris Chagnaud, and Parthena Schneider-Soupiadis for critical proofreading, and the Rupp lab and Hörmanseder lab for supply of *Xenopus* embryos. We further thank PD Dr. Steffen Dietzel and the center for advanced light microscopy at the LMU, as well as the center for advanced light imaging (CALM) for support and access to microscopes. This research was funded by the German Science Foundation (DFG, CRC 870, RTG 2175)

## Author contributions

HS and CG conceptualized, and HS, CG and MF designed experiments for the study. CG, MF and MK performed embryonic extirpations and quantified successful removals. CG and MK performed immunohistochemistry. MF and CG acquired behavioral data, and MF performed and imaged dye tracings. MF and CG established analysis scripts, and MF analyzed data. MF wrote the manuscript and created figures, and CG, MF edited the final manuscript. HS edited early versions of the manuscript.

## Data and code availability

All data reported in this paper will be shared by the <u>lead contact</u> on request. All code used for analysis of acquired behavioral data, as well as per-cycle segmented data, is publicly available on GitHub through Zenodo under http://doi.org/10.5281/zenodo.7781042.

## Material and Methods

### Animals

Experiments were conducted on wildtype *Xenopus laevis* embryos and tadpoles obtained from the animal breeding facility of the Biomedical Center at the Ludwig-Maximilians-University (LMU) Munich. Animals at early embryonic stages (stage 4-16) had their jelly coat removed with 2% cysteine and were incubated in 0.1x Marc’s modified Ringer solution (MMR, diluted from a 10x stock solution: 1 M NaCl, 18 mM KCl, 20 mM CaCl_2_, 10 mM MgCl_2_, 150 mM HEPES, pH 7.6–7.8) at 17°C under a 12/12 dark/light cycle regime. Animals were then reared until stage 46 either as control group, or with prior surgical removal of the embryonic optic vesicle at stage 26-28 (described below). At stage 46, a total of 83 manipulated animals were screened for successful removal of one eye (n = 22), and either euthanized with 3-aminobenzoic acid ethyl ester methanesulfonate (MS-222; Parmaq Ltd. UK) and fixed in 4% paraformaldehyde (PFA) for immunohistochemistry or were transferred into de-chlorinated water and further reared in standing tanks at 17-19°C at the animal facility of the Biocenter of the LMU Munich. Animals were kept on a 12/12 dark/light cycle, fed daily with powdered Spirulina (Algova, Germany) suspended in tank water, with a daily 50% water exchange. At stages 53-54, behavioral and anatomical experiments (described below) were performed on the animals in accordance with the “Principles of animal care” publication No. 86–23, revised 1985, of the National Institutes of Health and in accordance with the ARRIVE guidelines and regulations. Permission for the experiments was granted by the legally responsible governmental body of Upper Bavaria (Regierung von Oberbayern) under the license codes ROB-55.2.2532.Vet_03-17-24, ROB-55.2.2532.Vet_02-19-146 and ROB-55.2.2532.Vet_02-22-54. All experiments were performed in accordance with the relevant guidelines and regulations of the LMU Munich.

### Optic vesicle extirpation

Extirpation of the optic vesicle on the left side was performed at developmental Stages 26-28 (Fig. 1a, Suppl. vid. 1) with tungsten needles (0.125 mm, Fine Science Tools, 10130-05). Embryos were transferred into 1x MMR at 22°C and anesthetized with 0.02% Benzocaine (Sigma-Aldrich, E1501; see ^33^) prior to the surgical manipulations. The optic vesicle was visually identified, the overlaying ectoderm layer removed, and the optic vesicle excised along with the underlying optic stalk. Following this manipulation, animals were left in 1x MMR for 30 minutes to promote healing of the excision site and subsequently returned to 0.1x MMR for further rearing until stage 46. At this stage, animals were anesthetized in 0.02% benzocaine for visual inspection and sorted for successful eye removals under a stereo microscope (SteREO Discovery.V20, Axiocam 305 color camera, Carl Zeiss Microscopy GmbH). Successfully extirpated, one-eyed, tadpoles were then transferred to the animal facility for rearing until stage 53-54, when eye motion recordings were executed. Similar removals of sensory organs have been performed successfully, with no further detrimental effect on development^28,29,33^. A representative video (Suppl. Vid. 1) was taken with a color camera (Axiocam 405, Carl Zeiss Microscopy GmbH) under a stereo microscope (SteREO Discovery.V20, Carl Zeiss Microscopy GmbH) at 30 FPS (ZEN software 3.4.91, Carl Zeiss Microscopy GmbH).

### Immunohistochemistry

After sorting of phenotypes at stage 46, prototypic one-eyed tadpoles were prepared for the immunohistochemical analysis of nerve and muscle tissue around the site of the extirpated eye for further verification of a successful removal (Fig. 1B). To this end, animals were deeply anesthetized in 0.05% MS-222 and immersion-fixed in 4% PFA in 0.1x phosphate buffered saline (PBS, 0.3 mM Na_3_PO_4_, 15 mM NaCl, 0.105 mM K_3_PO_4_) for 3-6 hours at 4°C. Tadpoles were then transferred into 0.1x PBS, and lower jaws, viscera, tail, dorsal skull cartilage and brain were removed. The residual tissue was then dehydrated in 70% ethanol for 3-12 hours at 36°C, washed 3x in 0.1x PBS, and blocked in 5% normal goat serum in 0.1x PBS with 0.1% Triton X100 at 22-24°C for one hour. Incubation with the primary antibodies against muscle (MyosinVI, 1:400, Proteus Biosciences, 25-6791) and neuronal (acetylated tubulin, 1:800, Sigma-Aldrich, T7451) tissue was performed for 14 hours at 22-24°C. Subsequently, washing and blocking steps were performed as described above, followed by an incubation in secondary fluorescent antibodies (1:500, Alexa Goat anti-Mouse IgG2b, A-21141, Alexa Goat anti-Rabbit IgG, A32733) and DAPI (Thermo Fisher Scientific; 62248, 1:500) in 0.1x PBS with 0.1% Triton X100 for 1 hour at 22-24°C. Tissue was then washed 6 x for 15 minutes each in 0.1x PBS, mounted on microscope slides with spacers on each side, and coverslipped with Aqua Polymount (PolyScience, 18606). Imaging of the tissue was performed on a Leica SP5-2 confocal microscope (center for advanced light microscopy (CALM)).

### In vitro preparations

Activation and tracking of eye movements was performed in semi-intact *in vitro* preparations of either control or one-eyed tadpoles at stage 53-54. The generation of such isolated preparations occurred as described previously^56^. Animals were deeply anesthetized in 0.05 % MS-222 at 22-24°C for 2-5 minutes and were then transferred into ice-cold Ringer solution (75 mM NaCl, 25 mM NaHCO_3_, 2 mM CaCl_2_, 2 mM KCl, 0.1 mM MgCl_2_, and 11 mM glucose, pH 7.4). Following decapitation, the lower jaws, viscera and skin above the skull were removed, the skull opened and the choroid plexus above the IV^th^ ventricle was taken off. Preparations were then allowed to recover in 200 ml Ringer solution for ∼2 hours at 17°C before commencing with the eye motion recording session^33^. Unilateral optic nerve transections were performed directly before behavioral assessments. The optic nerve was located from dorsally through the opened skull and cut proximally to its exit point at the diencephalon with scissors under visual control.

### Visual motion (optokinetic) stimulation

Optokinetic stimuli consisted of horizontally moving vertical black and white bars (subtending 16° of visual angle each), projected with three orthogonally oriented digital light processing video projectors (Aiptek V60) onto a cylindrical screen covering 275° of the horizontal visual field of the tadpole. Stimulus motion consisted of either sinusoidal left-right oscillations or unidirectional constant velocity stimulus motion either to the left or to the right. Sinusoidal stimuli were presented at different frequencies (0.1 Hz, 0.2 Hz, 0.5 Hz; ±10° magnitude), presented in 2 trials of 15 cycles each, with a 30 second inter-trial interval. Constant velocity visual stimuli consisted of a 30 second stimulus motion at a velocity of 6.25°/s, with 30 seconds in either direction, with alternating starts to the left or to the right, with a stationary pattern for 30 seconds between the two motion directions. Stimulus motion was recorded and synchronized with the Spike2 software and a CED 1401 A/D interface (Cambridge Electronic Design, UK) at 50 Hz.

### Eye motion tracking

Preparations were recorded from above with a digital CCD camera (Grasshopper Mono, Point Grey Research Inc., Canada) equipped with an infrared high-pass filter and a zoom objective (Optem Zoom 70XL, Qioptiq Photonics GmbH & Co. KG, Germany) using infrared illumination of the preparation. Eye movements were tracked in real-time with a custom-written program^36^, measured as deflection angle between the vertical image axis and the long axis of an ellipse fitted around the two eyes which were detected by a manually adjusted darkness threshold. Eye positions were recorded in Spike2 at 30 Hz (Fig. 1c). In both control and one-eyed tadpoles, only data from the right eye was used for analysis.

### Retinal ganglion cell projections

Retinal ganglion cell (RGC) projections into the brain of control and one-eyed animals were fluorescently labelled following completion of the eye motion recording protocol with Tetramethylrhodamine, conjugated to 3000 MW Dextran (Invitrogen, D3308). The fluorescent dye was dissolved in ddH_2_O before crystallization onto 0.1 mm *minutiae* pins. *In vitro* preparations were mounted right lateral-side up, the cornea was incised and the lens removed. Following a transient removal of the Ringer solution and insertion of a needle with crystallized dye into the eye cup, the cornea was resealed with superglue (Uhu, Germany). Thereafter, *in vitro* preparations were incubated for 14-24 hours at 14°C and, following visual assessment of labelling quality and extent, immersion-fixed in 4% PFA at 4°C for 24 hours. Subsequently, the brain was extracted, DAPI-stained for 2 hours (1:500) and cleared with the uDISCO-Protocol^73^, with 2 hours per butanol step and clearing in BABB D-15. Cleared brains were mounted using custom built metal-spacers, sealed with Roti Histokitt II (Carl Roth, T160.1), and imaged on a Leica SP5-2 confocal microscope at an optical section spacing of 2.4 μm.

### Data analysis

#### Eye motion tracking

Eye movement data obtained in spike2 was processed as previously described^56^. Sinusoidal and unidirectional eye and stimulus motion data was converted into the matlab (Mathworks, USA) file format, non-uniformally sampled data resampled at 200 Hz and filtered with a 4 Hz low-pass filter. Sinusoidal eye movements were segmented into individual cycles, sorted for cycles that exclusively contained slow-following eye movements, and averaged. The amplitude and phase of the cyclic responses were measured based on magnitude and temporal occurrence of the response peaks, relative to the stimulus cycle. Eye movements evoked by constant velocity motion stimuli were similarly cut into individual episodes for each direction. The initial 0.75 seconds of each stimulus episode were discarded due to the response latency of motion onset. The following 2 seconds of each stimulus episode was then screened manually for resetting fast-phases. If fast-phases occurred prior to 1.5 second after stimulus onset, the episode was discarded; if fast-phases occurred after 1.5 second following stimulus onset, eye movements were included until fast-phase onset. Eye motion velocity was calculated as the mean derivative of the eye position and averaged across either leftward or rightward episodes per animal. In addition, an average leftward and rightward position trace was generated per animal for presentation purposes.

### Neuronal tracing

Image stacks acquired from cleared brains were loaded into FiJi as 8-bit images (Schindelin et al., 2012) and thresholded at an intensity of 3-4 depending on background fluorescence. Subsequently, a region-of-interest (ROI) was manually selected at the rostral boundary of the optic tectum across the whole medio-lateral extent of the brain and an intensity profile was generated for the ROI in each image of the stack. The ROI was limited to the caudal diencephalon and the rostral midbrain to measure intensity specifically in the approximate area of the pretectum. Intensity profiles were then summed up across the z-stack, normalized, and plotted, to generate a plot showcasing the amount of above-threshold labelled neuronal fibers within the ROI across the entire medio-lateral and dorso-ventral aspect of the brain. These plots were then split into left and right hemispherical halves, and the integral of each plot was calculated, approximating the total amount of fibers in the right and left (ipsi- and contralateral with respect to the right eye, respectively) rostral tectal region. The calculations served as proxy for the fraction of RGC axons projecting into the ipsi- and contralateral pretectum and optic tectum, respectively.

### Statistical analysis

Statistical comparisons between independent (one-eyed animals, control animals, left optic nerve-transected animals) for sinusoidal data were performed with a Kruskal-Wallis test for unpaired non-parametric data, followed by a Dunn’s multiple comparisons test to find group differences in Prism 9 (GraphPad Software Inc, USA). Statistical comparisons between and within animals (leftward/rightward eye movements for different groups) were tested with a 2-way ANOVA followed by a Bonferroni multiple comparisons test for group differences. The ROUT outlier test in Prism (maximum FDR = 0.1%) was performed to identify outliers in all plots, whit 1 data point removed from all plots in Figure 3 due to being an outlier in T/N asymmetry at a value of 2.36. To calculate motion direction preference for the left and right eye in control animals, preference of rightward/leftward eye movements for either unidirectional or oscillating eye movements, were calculated as the slope obtained by fitting the eye positions of both eyes plotted against each other, with the r^2^ indicating the quality of the fit (Python 3.7). Dots that are connected to each other with lines indicate paired data. Circular statistics were performed in Oriana (Version 4.02; Kovach Computing Services, see ^33^. A mean vector was calculated from phase values yielding the mean direction as well as the strength of the vector as an indication of data clustering (scale 0-1). Differences in phase values were identified with a Watson-Williams-F test.

## Abbreviations

N: nasal;
RGC: retinal ganglion cell;
T: temporal;
OKR: optokinetic reflex

## Supplemental material legends

*Supplemental Figure 1:*

Generation of one-eyed tadpoles. a) counts of animals generated from embryonic eye removals one week after extirpation. Monocular: successful generation of one-eyed animals with no residual eyes or adverse effects on surrounding tissue. Regrowth: regrowth of a partial or full eye, retina, or a lens. Deformations: Malformed tissue structures in the region of the extirpation or at the brain. b), c), Zoom-in into the eye region of a monocular (b) or control (c) tadpole at stage 46, 7 days after extirpations. Zoom-ins taken from images shown in Fig. 1b). d), e) *In-vitro* preparations of monocular (d) and control (e) tadpoles at stage 53, 4-5 Weeks after extirpations. Scalebars: b, c) 200 μm, d, e) 1000 μm

*Supplemental Figure 2:*

Left-right asymmetries in binocular and monocular tadpoles. a) Conjugations of the left and right eye for leftward (blue) and rightward (red) unidirectional motion, and for sinusoidal motion (black) in control animals (n=12). A slope of 1 represents conjugate eye motion, >1 stronger motion of the left, and <1 of the right eye. Slope of leftward and rightward motion are 1.12 ± 0.005 and 0.69 ± 0.005, respectively, while sinusoidal conjugation is intermediate at 0.92 ± 0.06. b) Conjugations of the left and right eye for leftward (blue) and rightward (red) components of sinusoidal motion. Slope of leftward and rightward motion are 0.95 ± 0.01 and 0.93 ± 0.01, respectively c) Plot taken from Fig. 3b on nasal and temporal gains during unidirectional stimuli of binocular and lesioned tadpoles. Added are grey dots which represent a summation of corresponding nasal and temporal values of lesioned animals (mean ± SD: 0.31 ± 0.08 p=0.08, two-way ANOVA, Bonferroni’s multiple comparison). d) Plot from Fig. 3c, with added grey dots which represent symmetry values of the lesioned nasal component, with the temporal component substituted with summed nasal and temporal gains calculated in panel (mean ± SD: 1.64 ± 0.54, p>0.99, Kruskal-Wallis test with Dunn’s multiple comparison. e) Correlation of nasal (red) and temporal (light blue) gains of embryonic monocular tadpoles (same individuals from Fig. 4f) with the amount of ipsilateral fraction of RGC projections. Nasal: R^2^ = 0.002, fit: y = -0.04y + 0.2. Temporal: R^2^ = 0.67, fit: y = -0.87y + 0.23

*Supplemental Video 1:*

Extirpation of the optic vesicle. Representative video of the removal of the left optic vesicle of a *Xenopus laevis* embryo at developmental Stages 26-28.

