## Supplementary figures and images for "Bilateral retinofugal pathfinding impairments limit behavioral compensation in near-congenital one-eyed *Xenopus laevis*"

### Supplemental figures

Supplemental figure 1

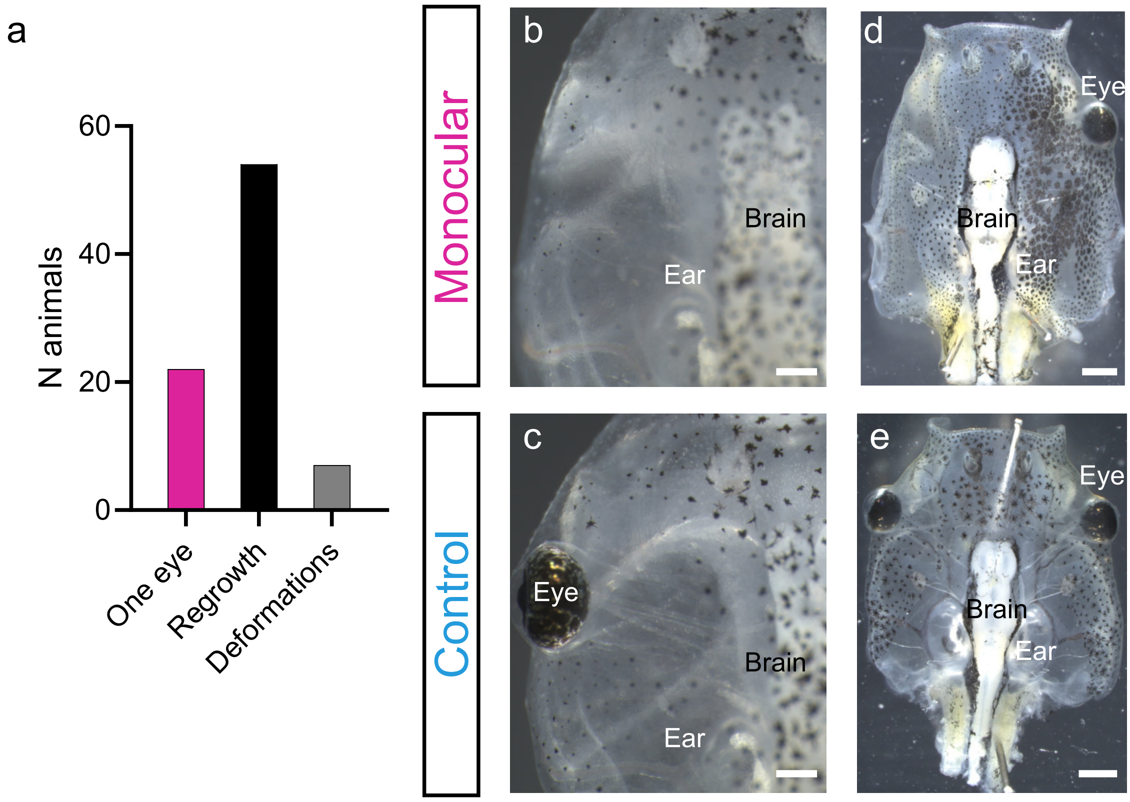

Supplemental figure 2

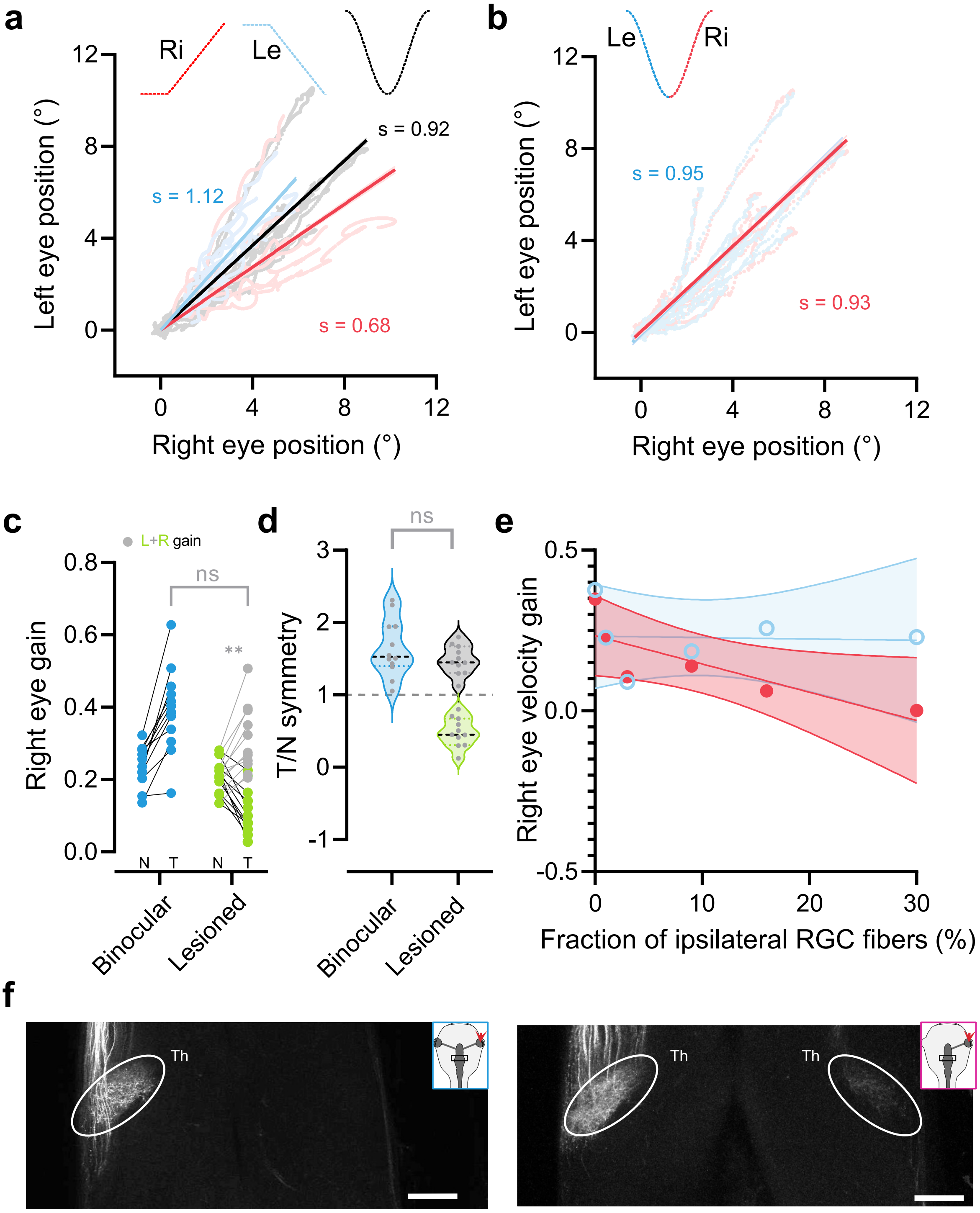
